# Loss of glutamate transporter *eaat2a* leads to aberrant neuronal excitability, recurrent epileptic seizures and hypoactivity

**DOI:** 10.1101/2021.03.16.435577

**Authors:** Adriana L. Hotz, Ahmed Jamali, Nicolas N. Rieser, Stephanie Niklaus, Ecem Aydin, Sverre Myren-Svelstad, Laetitia Lalla, Nathalie Jurisch-Yaksi, Emre Yaksi, Stephan C.F. Neuhauss

**Affiliations:** Department of Molecular Life Sciences, University of Zurich, CH-8057 Zurich, Switzerland; Life Science Zürich Graduate School, CH-8057 Zurich, Switzerland; Kavli Institute for Systems Neuroscience and Centre for Neural Computation, Faculty of Medicine and Health Sciences, Norwegian University of Science and Technology, 7030 Trondheim, Norway; Department of Neurology and Clinical Neurophysiology, St Olav University Hospital, 7030 Trondheim, Norway; Department of Neuromedicine and Movement Science, Faculty of Medicine and Health Sciences, Norwegian University of Science and Technology, 7030 Trondheim, Norway; Department of Clinical and Molecular Medicine, Norwegian University of Science and Technology, 7030 Trondheim, Norway; EraCal Therapeutics, CH-8952 Schlieren, Switzerland

## Abstract

Astroglial excitatory amino acid transporter 2 (EAAT2, GLT-1, SLC1A2) regulates the duration and extent of neuronal excitation by removing glutamate from the synaptic cleft. Hence, an impairment in EAAT2 function could lead to an imbalanced neural network excitability. Here, we investigated the functional alterations of neuronal and astroglial networks associated with the loss of function in the astroglia predominant *eaat2a* gene in zebrafish. We observed that *eaat2a*^-/-^ mutant zebrafish larvae display recurrent spontaneous and light-induced seizures in neurons and astroglia, which coincide with an abrupt increase in extracellular glutamate levels. In stark contrast to this hyperexcitability, basal neuronal and astroglial activity was surprisingly reduced in *eaat2a*^-/-^ mutant animals, which manifested in decreased overall locomotion. Our results reveal an essential and mechanistic contribution of EAAT2a in balancing brain excitability, and its direct link to epileptic seizures.

## INTRODUCTION

Astroglia are the most numerous glial cells within the central nervous system (CNS). They do not only provide trophic support to neurons but also play an important role in synapse formation and neurotransmission ^1–4^. By taking up neurotransmitters from the synaptic cleft, these glial cells are crucial for regulating synaptic transmission. The excitatory amino acid transporter 2 (EAAT2), expressed mainly on astroglia, plays a key role in synaptic regulation by removing the majority of extracellular glutamate, which is the main excitatory neurotransmitter in the CNS ^5,6^. Impaired glutamate clearance by this transporter has been shown to lead to synaptic accumulation of the neurotransmitter, resulting in an overactive CNS and excitotoxicity ^7,8^. This in turn may cause epilepsy, a group of brain disorders characterized by recurrent seizures ^9^.

Previous studies have shown that astroglia-neuron interactions may play an essential role in seizure initiation and propagation ^10–12^. On one hand, the functional coupling of astroglia through gap junctions is crucial to avert excessive neuronal activation, accomplished by rapid re-distribution of ions and neurotransmitters across the connected astroglial network ^12^. On the other hand, this functional syncytium might also, under special circumstances, promote epileptogenesis ^13^. The intercellular spread of calcium waves among astroglia can affect neuronal synchronization and therefore influence the propagation of seizure activity ^10^. In addition, disruptions of the glutamate-glutamine cycle, in which EAAT2 is essential, are linked with temporal lobe epilepsy in human patients and rodents ^14^. Accordingly, it is essential to understand how loss of EAAT2 affects neurons and astroglia.

In the present study, we generated a zebrafish (*Danio rerio*) mutant lacking EAAT2a, the zebrafish orthologue matching the mammalian EAAT2 in biophysical characteristics and mainly glial expression pattern ^15,16^. Calcium imaging in the transparent zebrafish larvae allowed us to image whole-brain network activity in vivo ^17^. We show that loss of EAAT2a transporter in larval zebrafish leads to increased brain excitability and recurrent spontaneous seizures, mimicking a human phenotype of patients with de novo mutations in EAAT2 ^18,19^. These seizures are manifested in zebrafish larvae by epileptic locomotor bursts and periods of excessive brain activity, accompanied by increased extracellular glutamate concentrations. Counterintuitively, apart from these periods of hyperexcitation, neuronal and astroglial network activity of *eaat2a*^-/-^ mutants is reduced. This coincides with a decreased overall locomotion compared to their unaffected siblings, and mirrors slow background brain activity and reduced muscle tone present in human patients ^18,19^. Altogether, our *in vivo* model of impaired EAAT2a function results in a depressed yet hyperexcitable brain state, and mimics a form of developmental and epileptic encephalopathy (DEE).

## RESULTS

### EAAT2a is predominantly expressed in astroglial cells

To investigate the expression pattern of *eaat2a* transcripts in the larval zebrafish, we first performed in situ hybridization experiments. Our results showed that *eaat2a* transcripts are expressed in all parts of the CNS (Fig. 1A). Specifically, we observed *eaat2a* transcripts along the spinal cord (Fig. 1A), the periventricular zones of the forebrain, the midbrain, and the hindbrain (Fig. 1B-D). In adult fish, the high expression in the forebrain (Fig. 1E) and tectal periventricular regions was maintained (Fig. 1F). These results suggest that spatial distribution of zebrafish *eaat2a* transcripts mainly overlaps with the location of astroglial cells, the functional homologues of mammalian astrocytes ^20,21^. To further confirm the precise expression of *eaat2a*, we performed triple staining using antibodies against astroglia and neuron specific proteins together with our custom made paralogue-specific antibody against EAAT2a ^16^ (Fig. 1G-L). We found substantial co-localization of EAAT2a staining with the astroglial glutamine synthetase (GS) along the ventricular zones (Fig. 1K), while neuronal cell labeling with SV2 (presynapse) and acetylated tubulin (axons) only showed weak EAAT2a signals (Fig. 1L). These results demonstrate that EAAT2a is mainly expressed in astroglial cells.

**Figure 1:**
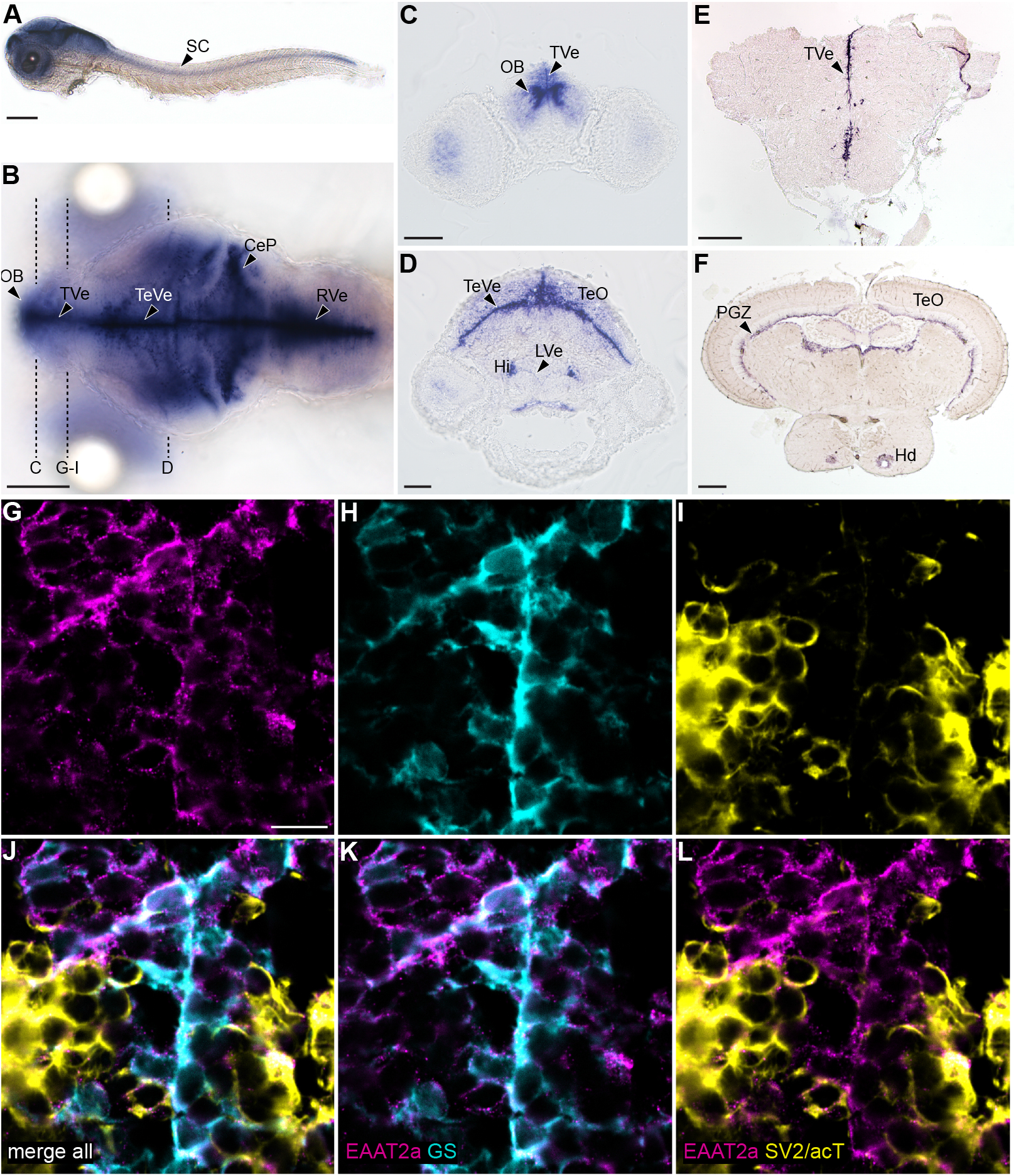
EAAT2a is predominantly expressed in astroglial cells. (**A** and **B**) mRNA of *eaat2a* is expressed along the periventricular zones (TVe, TeVe, RVe), resembling astroglial localization patterns in larval zebrafish as visible in lateral (**A**) and dorsal (**B**) view. (**C** and **D**) Cross sections of (**B**) indicated by dashed lines. (**E** and **F**) *eaat2a* mRNA expression along periventricular zones (TVe, PGZ) is maintained in adult zebrafish anterior forebrain (**E**) and midbrain (**F**). (**G-I**) Protein expression of EAAT2a (magenta), glutamine synthetase (GS, cyan) and synaptic vesicle 2/acetylated tubulin (SV2/acT, yellow) on cross sections of larval anterior forebrain indicated in (**B**). (**J-L**) Overlay of EAAT2a, GS and SV2/acT shows a greater co-localization of EAAT2a with astroglial (**K**) than neuronal (**L**) cells. CeP = cerebellar plate, Hi = intermediate hypothalamus, Hd = dorsal zone of periventricular hypothalamus, LVe = lateral ventricular recess of hypothalamus, OB = olfactory bulb nuclei, PGZ = periventricular gray zone of the optic tectum, TVe = periventricular zone of the telencephalon, TeO = optic tectum, TeVe = periventricular zone of the tectum, RVe = periventricular zone of the rhombencephalon. Scale bars are 200 μm in **A, B, E** & **F**; 50 μm in **C** and **D**; 10 μm in **G**.

### Loss of EAAT2a leads to morphological defects and larval lethality

To elucidate the function of EAAT2a in brain development and function, we generated CRISPR/ Cas9-mediated knockout (KO) mutants targeting exon 3 preceding the transmembrane domains involved in transport function ^22^. The selected mutant allele harbors a −13 base pair deletion, leading to premature STOP codons within the third transmembrane domain (Supplementary Fig. 1). The predicted truncated protein fragment is devoid of functional transport domains, consequently no EAAT2a antibody signal was present in *eaat2a*^-/-^ mutants (Supplementary Fig. 1 and 2). We observed that *eaat2a*^-/-^ zebrafish larvae displayed an aberrant morphology. They failed to inflate their swim bladder, developed pericardial edemas and were smaller than their siblings (Fig. 2A). Body size measurements confirmed that at 3, 4, 5, 6, and 7 days post fertilization (dpf), *eaat2a*^-/-^ zebrafish were significantly smaller than control animals (Fig. 2B), with reduced brain size at 5 dpf (Supplementary Fig. 3). Furthermore, *eaat2a*^-/-^ mutants were not viable past larval stage and started to die from 6 dpf on (Fig. 2C). Only 10% of *eaat2a*^-/-^ larvae were still alive at 9 dpf, and remaining survivors were heavily impaired, displaying large edemas and reduced locomotion. In contrast, *eaat2a*^+/-^ heterozygotes were indistinguishable from *eaat2a*^+/+^ siblings by visual inspection.

**Figure 2:**
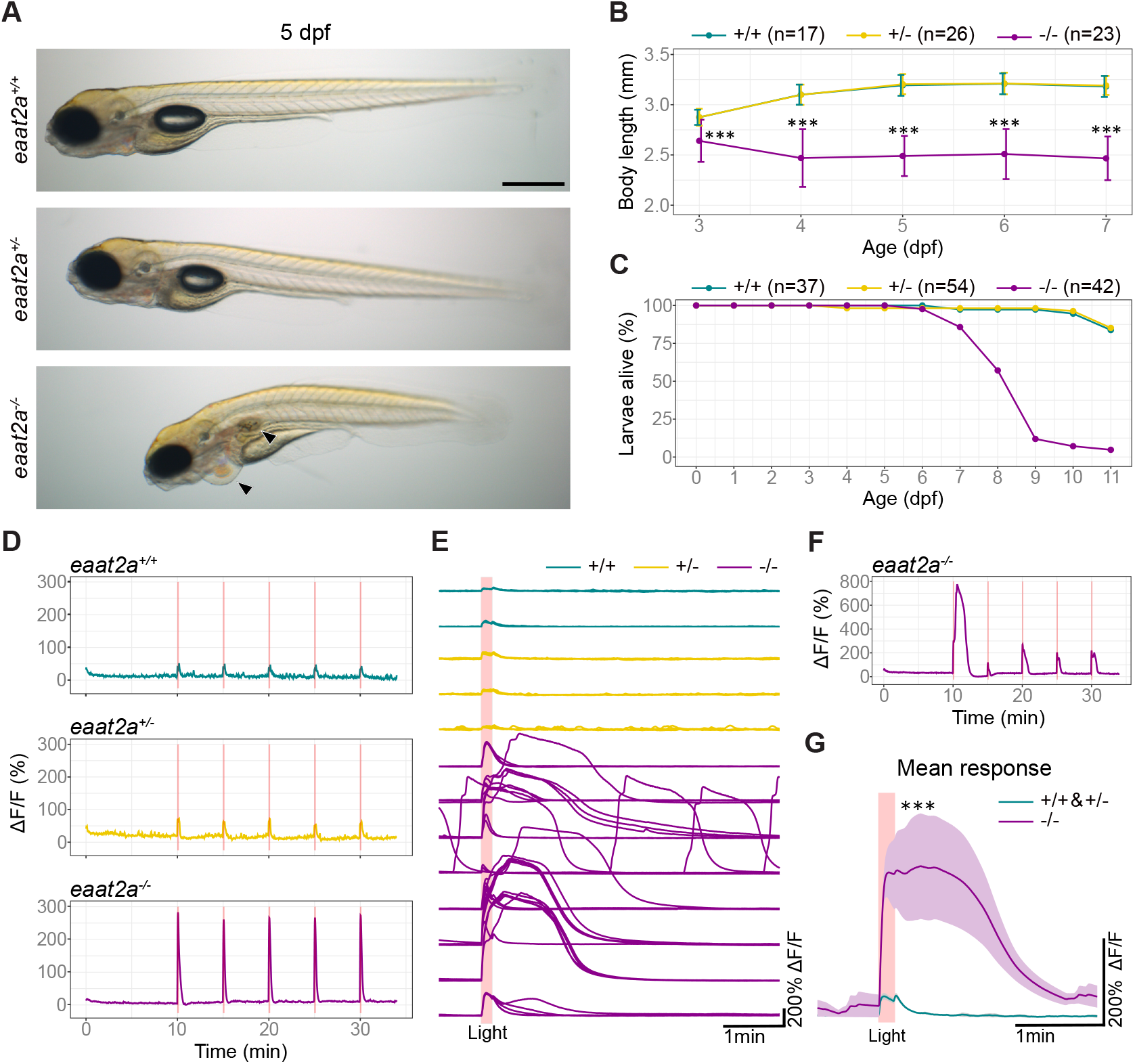
Knockout of *eaat2a* leads to hyperexcitability in response to light stimulation, morphological defects and larval lethality. (**A**) Lateral view of 5 dpf *eaat2a* mutants. *eaat2a*^-/-^ larvae (bottom) are slightly curved, do not have an inflated swim bladder and develop pericardial edema (arrowheads). *eaat2a*^+/+^ (top) and *eaat2a*^+/-^ (middle) larvae are indistinguishable. Scale bar is 500 μm. (**B**) Spinal cord length analysis of *eaat2a* mutants at consecutive days reveals a smaller body size in *eaat2a*^-/-^ (magenta) compared to their *eaat2a*^+/-^ (yellow) and *eaat2a*^+/+^ (cyan) siblings. (**C**) Survival curve of *eaat2a* mutants. (**D**) Whole-brain neuronal activity (*elavl3*:GCaMP6s signal) of three representative *eaat2a* mutant larvae (*eaat2a*^+/+^ in cyan, *eaat2a*^+/-^ in yellow and *eaat2a*^-/-^ in magenta) exposed to five 10-second light stimuli with 5-minute interstimulus interval. (**E**) Changes in fluorescence over time (ΔF/F) per larvae aligned at the onset of light stimuli. (**F**) ΔF/F trace of an example *eaat2a*^-/-^ larvae with diverse responses to light-stimuli. (**G**) Mean responses to light stimuli in 5 dpf *eaat2a*^+/+^ and *eaat2a*^+/-^ (cyan), and *eaat2a*^-/-^ (magenta) zebrafish larvae. Shaded area represents SEM. Significance level: \*\*\**p* = < 0.001, Kruskal Wallis test with Wilcoxon rank-sum post hoc test (**B**) or Wilcoxon rank-sum test (**G**).

### *eaat2a*^-/-^ mutant zebrafish exhibit hyperexcitability in response to light stimulation

EAAT2 is an essential part of the glutamate clearance mechanism in the brain ^5,16^. Hence, the absence of the glutamate transporter may lead to changes in neuronal excitability in *eaat2a*^-/-^ mutant zebrafish. To test this hypothesis, we compared brain-wide neuronal responses to transient light flashes in *eaat2a*^-/-^ zebrafish and control siblings expressing the transgenic calcium indicator GCaMP6s under the neuronal *elavl3* promoter ^10^. We observed that *eaat2a*^-/-^ mutant larvae displayed highly amplified light responses compared to their *eaat2a*^+/-^ and *eaat2a*^+/+^ siblings (Fig. 2D and E), with varying response amplitudes across and within *eaat2a*^-/-^ larvae (Fig. 2F). Strikingly, average neuronal responses in *eaat2a*^-/-^ larvae were not only excessively enlarged, but also very long lasting (Fig. 2G). Occasionally, we also observed spontaneous neuronal activity bursts of similar magnitude (Fig. 2E), potentially resembling spontaneous epileptic seizures ^9,23-25^. Taken together, our results show that *eaat2a*^-/-^ zebrafish brains exhibit increased excitability in responses to light stimulation and are possibly prone to epileptic seizures.

### *eaat2a*^-/-^ mutant zebrafish show spontaneous seizures coinciding with a surge of extracellular glutamate

Our neuronal activity recordings suggested that *eaat2a*^-/-^ mutant zebrafish not only exhibit strong light-induced responses but also display occasional neuronal activity bursts resembling epileptic seizures. To further characterize this spontaneous seizure-like phenotype in *eaat2a*^-/-^ mutants, we examined swimming behavior in 5 dpf larvae by using automated behavioral tracking. We observed that *eaat2a*^+/+^ and *eaat2a*^+/-^ zebrafish swim with periods of stop, slow and fast swims as described previously in healthy zebrafish ^26,27^. In contrast, *eaat2a*^-/-^ mutant animals displayed aberrant locomotor patterns (Fig. 3A-F). *eaat2a*^-/-^ larvae swam substantially less than their *eaat2a*^+/-^ and *eaat2a*^+/+^ siblings, mainly lying motionless on the bottom of the dish (Fig 3A-C). Strikingly, when *eaat2a*^-/-^ larvae swam, they showed convulsive twitching and swim bursts occasionally followed by swirling around their body axis and finally immobilized sinking to the bottom of the dish (Supplementary Movie 1). Compared to their siblings, *eaat2a*^-/-^ zebrafish showed more swim bursts of either long distance (Fig. 3D and E) or high velocity during a prolonged period (Fig. 3F, Supplementary Fig. 4). This swirling and bursting behavior highly resembles established models of epilepsy and is interpreted as seizure-like behavior in zebrafish ^24,28^. All these findings support the idea that *eaat2a*^-/-^ larvae exhibit spontaneous seizures.

**Figure 3:**
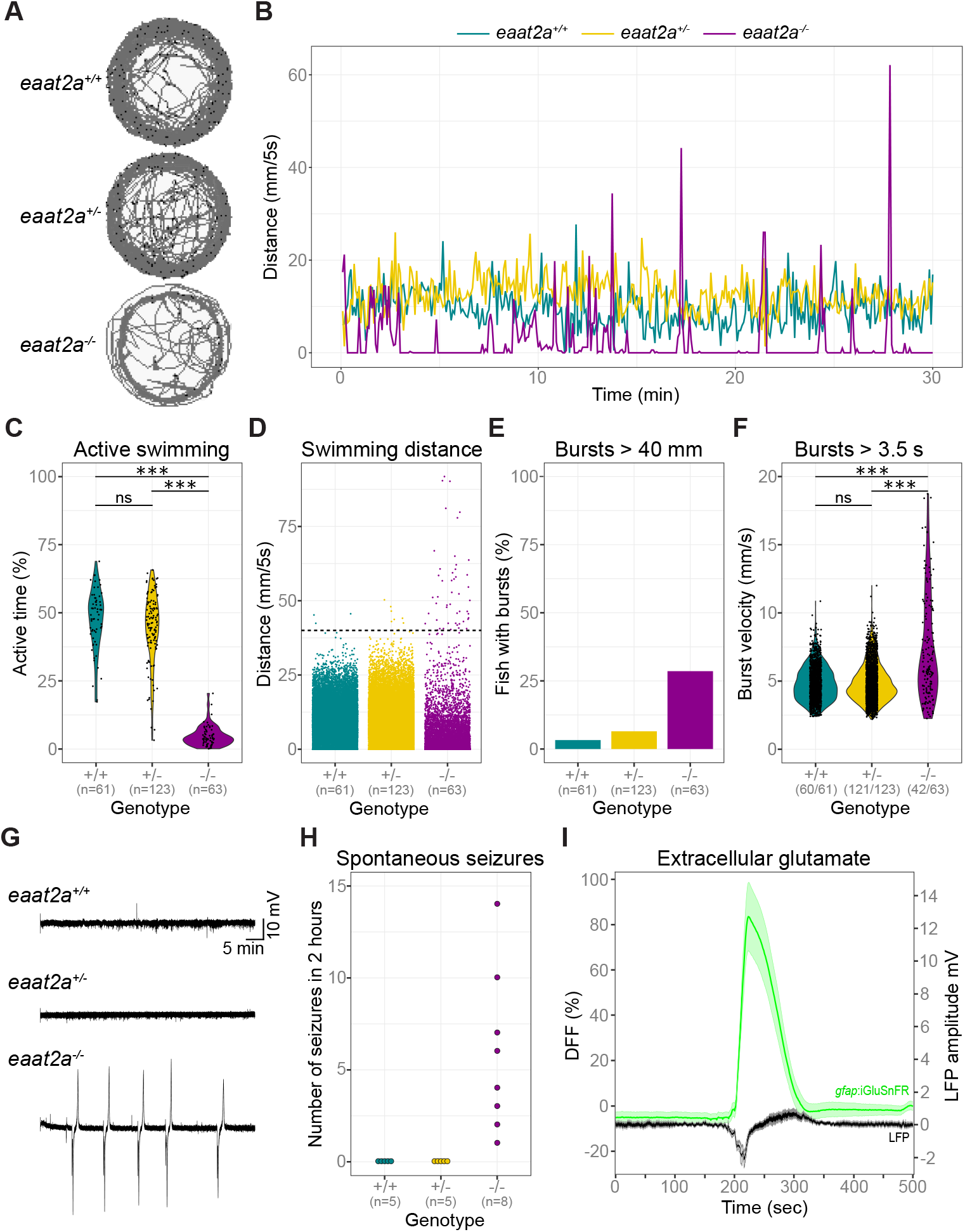
*eaat2a*^-/-^ mutants show spontaneous seizures coinciding with a surge of extracellular glutamate. (**A** and **B**) Representative traces for swim location (**A**) and distances (**B**) over 30 minutes in 5 dpf *eaat2a*^-/-^ (magenta), *eaat2a*^+/-^ (yellow) and *eaat2a*^+/+^ (cyan) larvae. (**C**) Ratio of time period with active swimming, per genotype. (**D**) Distance moved per five-second integral during 30-minute recordings, per genotype. Black dotted line represents threshold > 40 mm/5 s. (**E**) Proportion of fish showing one or more bursts bigger than 40 mm during five-second integrals. (**F**) Velocity of all bursts lasting longer than 3.5 seconds. (**G**) Representative telencephalic local field potential (LFP) recordings. (**H**)Number of spontaneous global seizures detected during two-hour LFP recordings. (**I**) Average change in iGluSnFR fluorescence (ΔF/F, green) and simultaneous local field potential (LFP, black) recordings during epileptic activity in 5 dpf *eaat2a*^-/-^ mutants. n = 8 fish. Signals are aligned at the onset of global seizures. Significance levels: \*\*\**p* = < 0.001, ns = not significant (*p* > 0.05), Kruskal-Wallis rank-sum test with Wilcoxon rank-sum posthoc test (**C**) or two-sample Kolmogorov-Smirnov test (**F**).

To verify the presence of seizure-activity in *eaat2a*^-/-^ mutant brains, we measured local field potentials (LFPs) by inserting a micro-electrode in the anterior forebrain (telencephalon) ^10^ of 5 dpf animals. As expected, the measured electrical activity in *eaat2a*^-/-^ larvae revealed spontaneous episodes of high voltage LFP deflections (Fig. 3G), resembling seizure-like LFP activity in other zebrafish models ^10,29,30^. These spontaneous seizures were not detected in *eaat2a*^+/-^ or *eaat2a*^+/+^ siblings (Fig. 3G and H). We hypothesized that spontaneous seizures in *eaat2a*^-/-^ mutant larvae are associated with an excess of extracellular glutamate that cannot be removed due to impaired glutamate clearance ^31^. Hence, we expected to see large glutamate surges during spontaneous seizures. To test this hypothesis, we combined LFP measurements with simultaneous fluorescence recordings of extracellular glutamate near astroglial terminals using *Tg(gfap:iGluSnFR)* animals ^32^. In *eaat2a*^-/-^ larvae, we observed massive increases of iGluSnFR signal reflecting glutamate levels, coinciding with spontaneous LFP-seizures (Fig. 3I). No such large glutamate surges were observed in *eaat2a*^+/-^ and *eaat2a*^+/+^ control siblings. Taken together, our results show that loss of EAAT2a leads to spontaneous seizures that coincide with a massive surge in extracellular glutamate levels.

### Neuronal hyperactivity during seizures contrasts with basal hypoactivity in *eaat2a*^-/-^ larvae

To characterize EAAT2a-related seizures further, we investigated the neuronal activity in 5 dpf *Tg(elavl3:GCaMP5G)* zebrafish. Our recordings revealed recurrent periods of excessive neuronal activity spreading across the entire brain of *eaat2a*^-/-^ mutants (Fig. 4A, Supplementary Movie 2 and 3). During these spontaneous seizures, neuronal calcium signals across the brain reached levels greater than 100% of relative change in fluorescence (ΔF/F) (Fig. 4B bottom). These globally high levels of neuronal seizure-activity were maintained for more than a minute before decreasing to a short hypoactive period, and finally returning to inter-ictal (between seizures) levels (Fig. 4B bottom and 4C). These results confirm that *eaat2a*^-/-^ mutants exhibit spontaneous global seizures that are not present in *eaat2a*^+/-^ and *eaat2a*^+/+^ control siblings (Fig. 4D). In addition to these global seizures, we found that some larvae additionally or exclusively showed localized seizures not reaching the anterior forebrain and lasting for less than a minute (Supplementary Fig. 5). Next, we asked how different brain regions are recruited during seizure propagation in *eaat2a*^-/-^ mutants. As represented in Figure 4E, excessive increase in intracellular calcium levels was initiated in the midbrain (8 of 12 fish) or hindbrain (4 of 12 fish). To quantify this further, we compared half maxima of neuronal calcium signals (ΔF/F) between anterior forebrain, midbrain and hindbrain during global seizures lasting for more than one minute. We observed that neurons of the anterior forebrain were recruited only seconds after seizure initiation (mean 13 sec, SD ± 5.67 sec), regardless of the seizure origin (Fig. 4F). Beyond these spontaneous seizures, we also observed that the large amplitude light responses shown earlier in Fig. 2E were very similar to spontaneous global (Fig. 4G, curly brackets) and localized (Fig. 4G, square brackets) seizures with comparable amplitudes (Fig. 4H). Since epileptic seizures that are objectively and consistently evoked by a specific external stimulus are referred to as reflex seizures, we termed these excessive responses to light in *eaat2a*^-/-^ mutants as light-induced reflex seizures ^33^. Intriguingly, during inter-ictal periods, *eaat2a*^-/-^ mutants exhibited reduced basal neuronal activity compared to *eaat2a*^+/-^ and *eaat2a*^+/+^ siblings (Fig. 4I). We observed smaller fluctuations (standard deviation) of neuronal calcium signals in *eaat2a*^-/-^ larvae compared to their siblings (Fig. 4J). This reduced basal activity might explain hypoactive locomotor behaviors of *eaat2a*^-/-^ mutants (Fig. 3C). Taken together, our results indicate that loss of EAAT2a not only leads to spontaneous seizures, but also reduces basal neuronal activity during inter-ictal periods. Moreover, in all our experiments, *eaat2a*^+/-^ mutants were indistinguishable from their *eaat2a*^+/+^ siblings, suggesting that *eaat2a*^+/-^ mutation does not lead to haploinsufficiency or any seizure phenotype.

**Figure 4:**
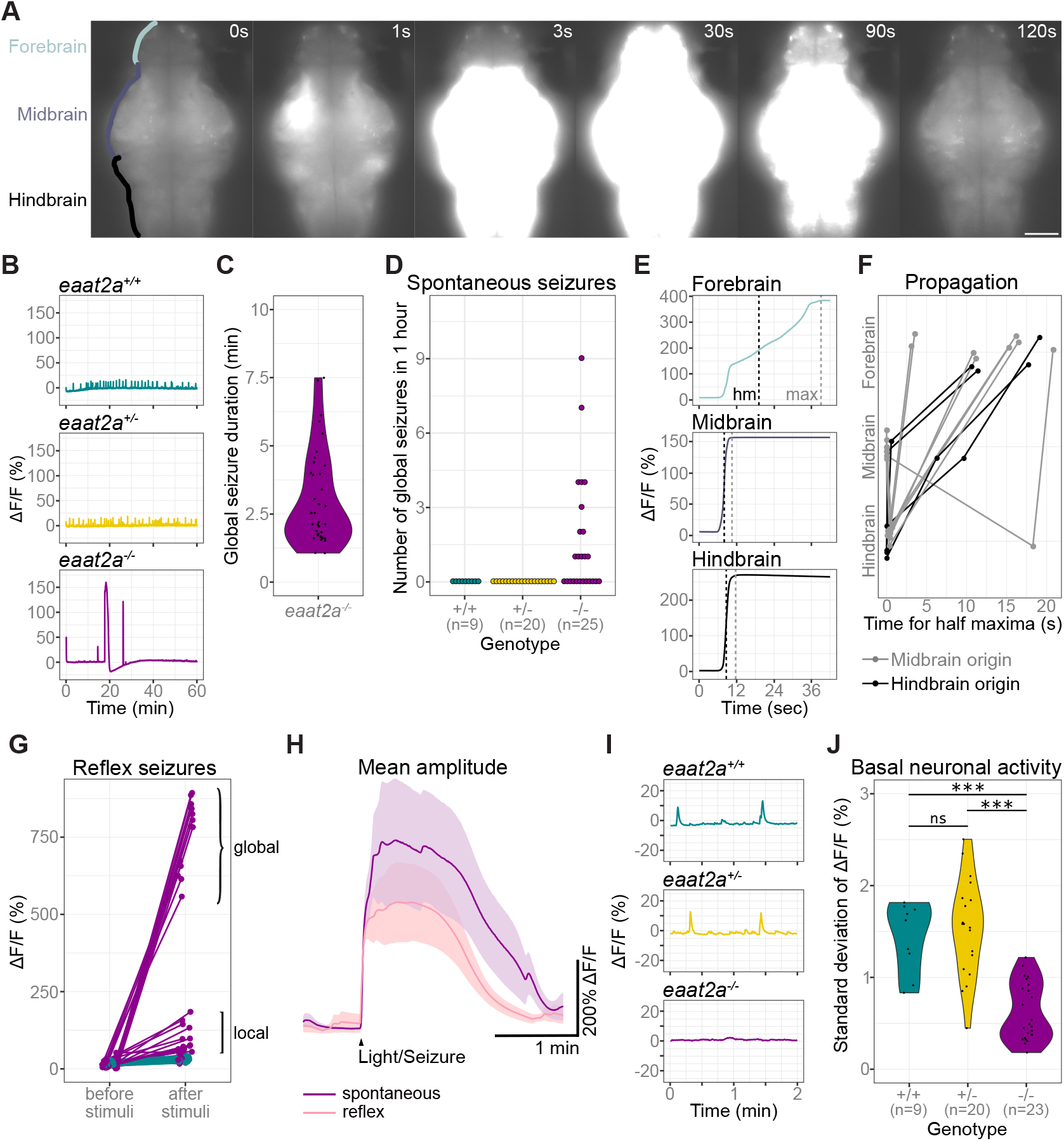
Neuronal hyperactivity during seizures contrasts with basal hypoactivity in *eaat2a*^-/-^ larvae. (**A**) Time lapse of neuronal calcium signals during a seizure in *eaat2a*^-/-^ larva in *Tg(elavl3:GCaMP5G)* background (dorsal view). Scale bar is 100 μm. (**B**) Representative calcium signals *(elavl3:GCaMP5G)* recorded across the brain of *eaat2a*^+/+^ (cyan, top), *eaat2a*^+/-^ (yellow, middle) and *eaat2a*^-/-^ (magenta, bottom) larvae. (**C**) Duration of spontaneous global seizures present in *eaat2a*^-/-^ mutants (median 2 min 7 sec (std ± 52 sec)). (**D**) Number of spontaneous global seizures recorded by calcium imaging during 60 minutes. (**E**) Neuronal activity *(elavl3:GCaMP5G)* of the three main brain regions of a representative global seizure. The half maxima (hm) represents the time point of max(ΔF/F)/2. (**F**) Relative time for ΔF/F half maxima represents propagation of global seizures across brain parts over time. Colors indicate region of seizure origin: light grey for midbrain, black for hindbrain. n = 12 *eaat2a*^-/-^ larvae. (**G**) Calcium signals (*elavl3*:GCaMP6s) one minute before and immediately after light stimuli. Brackets indicate global (curly brackets) and local (square brackets) reflex seizures. (**H**) Averaged calcium signals for spontaneous (magenta) and reflex (light pink) seizures during light-stimuli recordings. Shaded area represents SEM. (**I**) Magnification of (**B**) shows calcium signals (*elavl3:GCaMP5G*) during two-minute basal activity period used for standard deviation calculations in (**J**). (**J**) Neuronal basal activity calculated by the standard deviation of ΔF/F over two minutes in *eaat2a*^-/-^ mutants (mean 0.63, n = 23) compared to their *eaat2a*^+/-^ (mean 1.54, n = 20, *p* = 2.7e-07) and *eaat2a*^+/+^ (mean 1.44, n = 9, *p* = 1e-04) siblings. Significance levels: \*\*\**p* = < 0.001, ns = not significant (*p* => 0.05), Kruskal Wallis test with Wilcoxon rank-sum posthoc test (**I**).

### Astroglial network in *eaat2a*^-/-^ mutants is silent yet hyperexcitable

Given the predominant astroglial expression of EAAT2a, one potential cause for the reduced neuronal basal activity in *eaat2a*^-/-^ mutants may be the impaired astroglial glutamate recycling, which reduces available glutamate. To investigate whether astroglial function is impaired in *eaat2a*^-/-^ mutants, we recorded glial calcium signals in 5 dpf larvae expressing GCaMP6s under the glial promoter *gfap(Tg(gfap:Gal4)nw7;Tg(UAS:GCaMP6s)* ^10^. To exclusively analyze astroglial activity, we focused on the region along the ventricular zones in the forebrain and midbrain (Fig. 5A) ^10^. In line with our previous results, we observed spontaneous events of excessive astroglial activity (ΔF/F greater than 100 %) spreading across the entire brain of *eaat2a*^-/-^ mutants (Fig. 5B), often finally recruiting the anterior forebrain (Fig. 5C, Supplementary Movie 4 and 5). These events likely represent astroglial activity during seizures ^10^. Such spontaneous global events were prominent at least once per hour in *eaat2a*^-/-^ mutants and absent in *eaat2a*^+/-^ and *eaat2a*^+/+^ control siblings (Fig. 5D). To further characterize astroglial activity, we quantified the ratio of active glial cells and the amplitudes per calcium burst in *eaat2a*^-/-^ mutant larvae during inter-ictal, pre-ictal (preceding seizure onset) and ictal periods. We observed a drastic increase in the ratio of active astroglia and amplitude of calcium bursts at the transition from pre-ictal to ictal period, while inter-ictal and pre-ictal periods appeared to be similar (Fig. 5E and F). Next, we tested whether these large bursts of astroglial calcium signals are also present in light-evoked seizures. Our results revealed that the ratio and the amplitude of astroglial calcium signals were significantly larger in *eaat2a*^-/-^ mutants, when compared to control siblings (Fig. 5G and H). Furthermore, some of these light-evoked seizures propagated globally, recruiting the entire brain including the anterior forebrain (Fig. 5H, arrowheads). During inter-ictal periods, a smaller ratio of astroglial cells were active (Fig. 5I), and these active cells had reduced overall activity (Fig. 5J) in *eaat2a*^-/-^ mutants compared to their siblings’. However, we did not observe a significant difference in the amplitude or the duration of individual astroglial calcium bursts between *eaat2a*^-/-^ mutants and control siblings during inter-ictal/basal periods (Fig. 5K and L). Taken together, our results revealed that astroglial cells in *eaat2a*^-/-^ mutants are less active during inter-ictal states, and generate excessive calcium bursts only during seizures.

**Figure 5:**
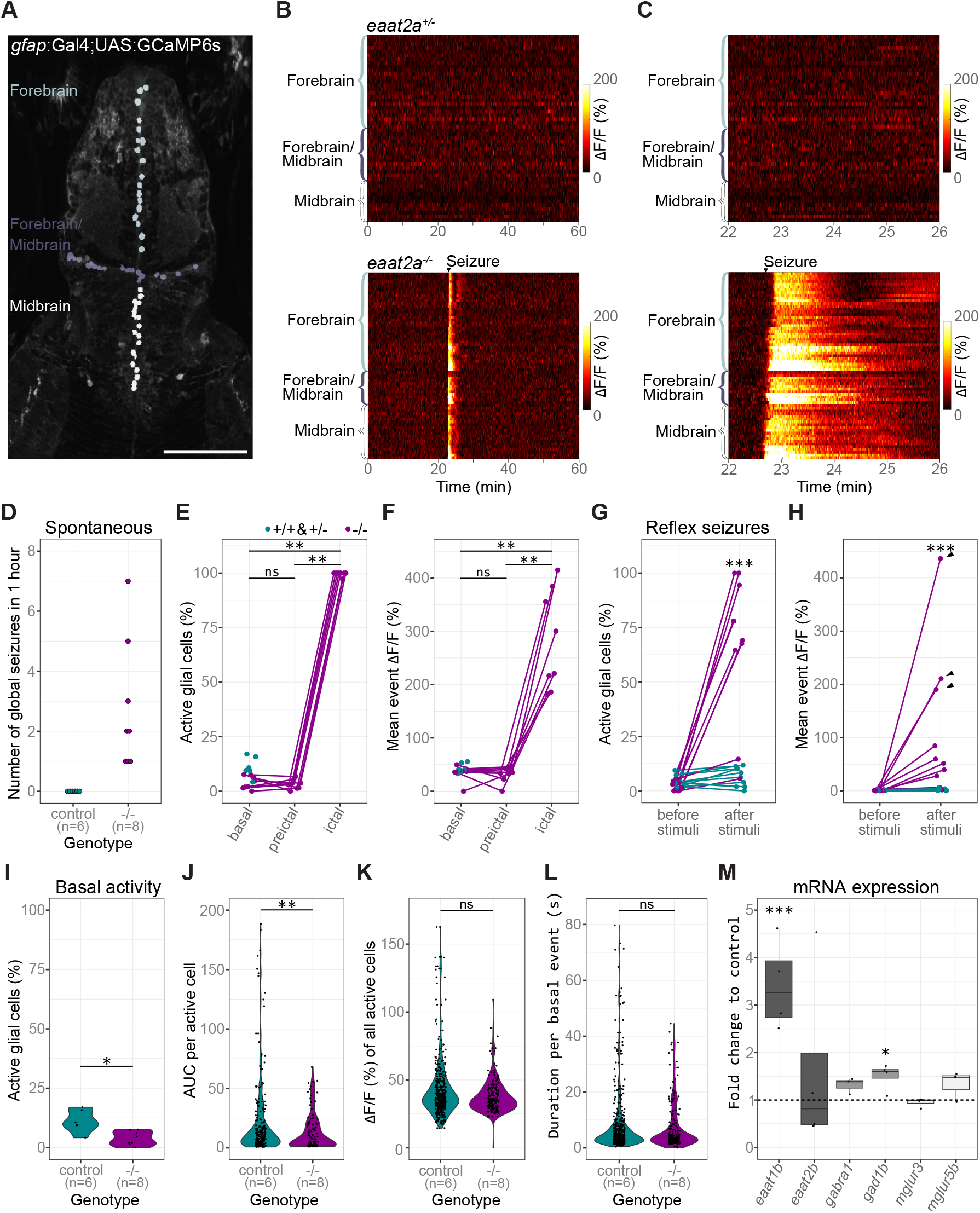
Astroglial network in *eaat2a*^-/-^ mutants is silent yet hyperexcitable. (**A**) Two-photon microscopy image of the forebrain and midbrain in a 5 dpf *Tg(gfap:Gal4;UAS:GCaMP6s)* zebrafish larva expressing GCaMP6s in GFAP positive glial cells. Individual astroglia along the ventricular regions are color-coded according to three different areas: forebrain (light grey), forebrain/mid-brain boundary region (dark grey), and midbrain (white). Scale bar is 100 μm. (**B**) Calcium signals (ΔF/F) of individual glial cells over time in representative *eaat2a*^+/-^ (top) and *eaat2a*^-/-^ (bottom) larvae. Warm color indicates high activity as seen during the seizure in the *eaat2a*^-/-^ larva. (**C**) Four-minute periods of calcium signals in (**B**). (**D**) Number of global seizures detected in *eaat2a*^-/-^ mutants (magenta) compared to their *eaat2a*^+/-^ and *eaat2a*^+/+^ siblings (cyan). (**E** and **F**) Percentage of active glial cells (**E**) and average event amplitudes of active cells (**F**) of individual *eaat2a*^-/-^ mutants during two-minute basal, preictal and ictal (spontaneous seizures) periods (**E**: *p_b-p_* = 1, *p_b-i_* = 0.0078, *p_p-i_* = 0.0078; **f**: *p_b-p_* = 0.95, *p_b-i_* = 0.0078, *p_p-i_* = 0.0078). Cyan dots show average basal values of *eaat2a*^+/+^ and *eaat2a*^+/-^ siblings. (**G** and **H**) Evaluation of one-minute periods immediately before and after 10-second light stimuli in *eaat2a*^+/+^/*eaat2a*^+/-^ (cyan) and *eaat2a*^-/-^ (magenta) animals of the proportion of active cells (**G**, *p* = 8.7e-4) and averaged amplitude over all cells (**H**, *p* = 6.7e-4). Arrowheads indicate global seizures. n = 5 fish per group, 2 stimuli per fish. (I**-L**) Analysis during inter-ictal basal periods of percentage active astroglia per fish (**I**, *p* = 0.011), total activity of active cells (**J**, area under the curve = AUC, *p* = 0.0011), amplitudes (**K**, *p* = 0.052) and durations (**L**, *p* = 0.37) of individual calcium bursts in active glial cells are plotted in violin plots with individual data points (black). n(*eaat2a*^+/+^/*eaat2a*^+/-^) = 6, n(*eaat2a*^-/-^) = 8. (**M**) mRNA transcript levels of *eaat1b* (*p* = 0.00075), *eaat2b* (*p* = 0.94), *gabra1* (*p* = 0.40), *gad1b* (*p* = 0.024), *mglur3* (*p* = 0.86) and *mglur5b* (*p* = 0.40) in 5 dpf *eaat2a*^-/-^ relative to *eaat2a*^+/+^ siblings. Transcripts were measured by RT-qPCR and normalized to *g6pd* and *b2m*. Significance levels: \*\*\**p* = < 0.001, \*\**p* = < 0.01, \**p* = < 0.05, ns = not significant (*p* => 0.05), Wilcoxon signed rank test (**E, F**), Welch two sample unpaired t-test (**I, M**) or Wilcoxon rank-sum test (**G, H, J-L**).

The depressed yet hyperexcitable brain state of *eaat2a*^-/-^ mutants implies pathological changes in the brain. To test the impact of *eaat2a* loss on the gene expression of key regulators of neuronal and astroglial activity, we analyzed whole-brain transcript levels using quantitative reverse transcription PCR (RT-qPCR). Firstly, we tested for transcriptional adaptation of other *eaat* glutamate transporters. Expression levels of *eaat2b*, the *eaat2a* paralogue, did not differ in *eaat2a*^-/-^ animals (Fig. 5M). In contrast, another glutamate transporter *eaat1b* was upregulated in *eaat2a*^-/-^ mutants compared to *eaat2a*^+/+^ siblings (Fig. 5M), suggesting a compensatory effect to improve buffering of excess synaptic glutamate in *eaat2a*^-/-^ mutants (Fig. 3I). Secondly, we assessed whether expressions of genes associated with inhibitory neurotransmission of γ-aminobutyric acid (GABA) are altered. We found that neither the expression of the GABA_A_ receptor subunit *gabra1* nor the presynaptic enzyme *gad1b* differed considerably between *eaat2a*^-/-^ and *eaat2a*^+/+^ brains (Fig. 5M). Finally, we investigated whether metabotropic glutamate receptors (mGluRs) on astroglia are affected. These receptors can influence intracellular calcium transients in astroglia following synaptic glutamate release ^3^. In *eaat2a*^-/-^ larval brains, we observed no difference in expression levels of glial expressed *mglur3* or brain abundant ^34^ *mglur5b*. This suggests that reduced inter-ictal activity in *eaat2a*^-/-^ mutant astroglia is not due to impaired mGluR-induced calcium signaling. Taken together, the altered gene expression levels of *eaat1b* in *eaat2a*^-/-^ mutants suggest a compensatory attempt to adjust increased brain excitability by buffering glutamate.

## DISCUSSION

In sum, we show that glutamate transporter EAAT2a is required to balance brain excitability by regulating extracellular glutamate levels. Our results indicate that impaired EAAT2a function results in epileptic seizures but also reduced basal brain activity (Fig. 6). Epileptic seizures are due to an imbalance between excitatory and inhibitory synaptic transmission. In this study, we show that a genetic perturbation of the excitatory glutamate system leads to profound functional alterations in neuronal and astroglial networks, leading to spontaneous and light-induced seizures. Our results are in line with the earlier observations associating EAAT2 malfunctions to seizures in mice and humans ^18,19,35–39^. In addition, our novel *eaat2a*^-/-^ mutant zebrafish epilepsy model has several important characteristics. Firstly, homozygous mutations of the astroglial *eaat2a* in zebrafish resemble a form of DEE present in human epilepsy patients with de novo mutations in the orthologous gene (EAAT2 = SLC1A2)^18,19,37^. DEE is clinically defined as a condition where developmental defects due to a mutation as well as epileptic activity itself contribute to the impairments^40^. The presence of both recurrent spontaneous seizures and severe developmental abnormalities in *eaat2a*^-/-^ mutants mirror the human phenotype ^18,19^. Secondly, spontaneous recurrent seizures in our *eaat2a*^-/-^ mutants enable the investigation of epilepto-genesis throughout development. Although *eaat2a*^-/-^ mutants’ premature death restricts analysis to larval stages, future work on an inducible gene knockout would allow studies at more advanced developmental stages ^41^. Finally, knockout of EAAT2a directly targets mainly astroglial networks, which are of great importance during seizure initiation and propagation ^10–12^. Therefore, our model will enable further investigations into the role of glia-neuron interactions in epilepsy. All these points give a novel vantage point for modelling and understanding potential mechanisms underlying human genetic epilepsies.

**Figure 6:**
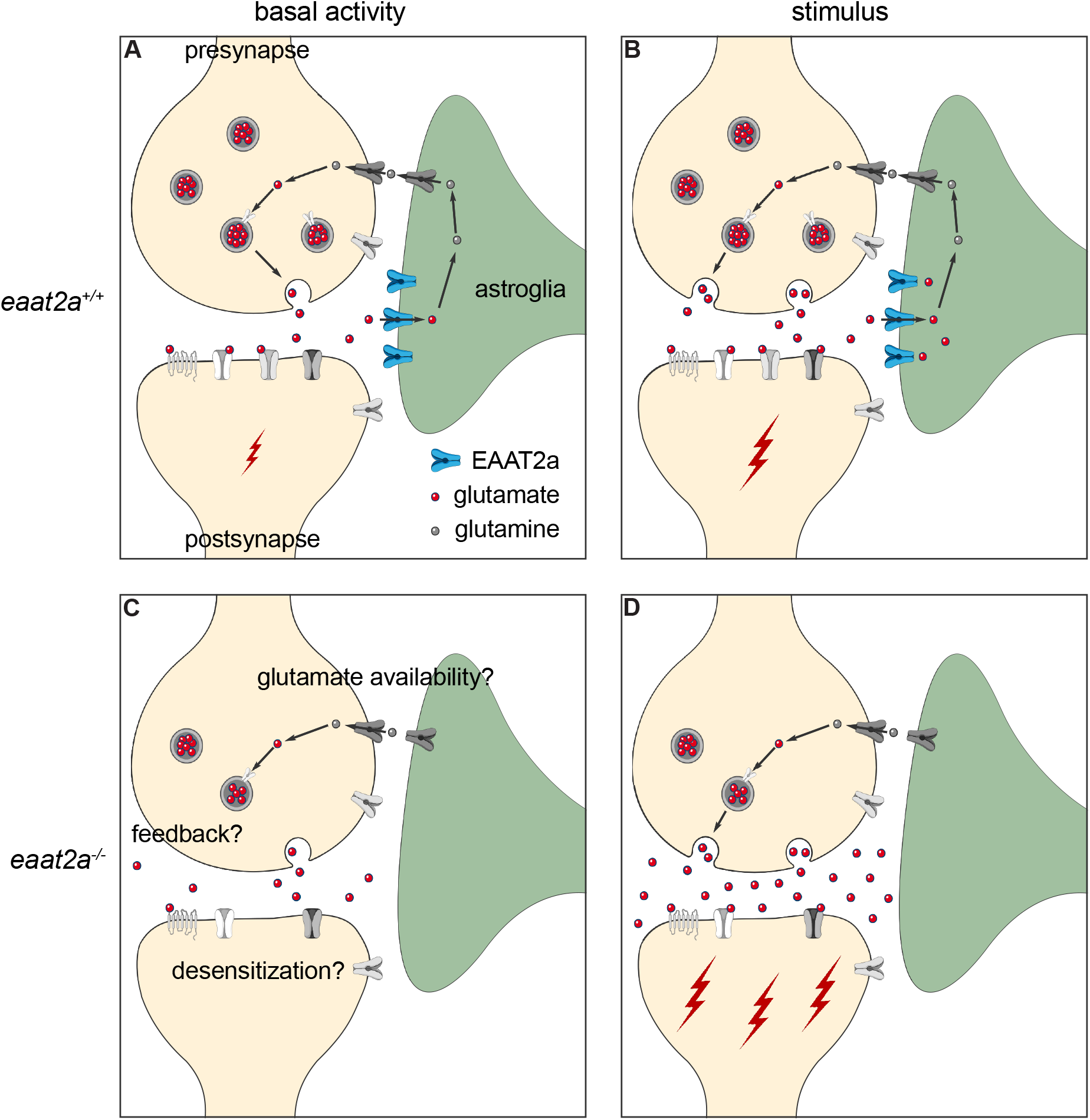
Working model of coexisting hyper- and hypoactivity in *eaat2a*^-/-^ mutants. (**A** and **B**) Glutamate transporter EAAT2a is important for both clearance of glutamate at the synaptic cleft and recycling of glutamate to the presynapse during basal activity and stimulation. (**C**) Loss of EAAT2a function leads to neuronal hypoactivity during basal periods, likely reflecting reduced levels of available glutamate/glutamine. (**D**) Once glutamate is released in higher quantities from the presynapse (e.g. following light stimuli or spontaneous glutamate release), astroglia cannot sufficiently take up glutamate. Accumulated glutamate in the synaptic cleft hyperexcites postsynaptic neurons, which can lead to a cascade of glutamate release across the nervous system and results in epileptic seizures. Such recurrent epileptic seizures could also potentially lead to desensitization or a negative feedback to the presynapse (**C**), which can further explain reduced basal neuronal activity in *eaat2a*^-/-^ mutants.

Recent studies show that astroglia and their interactions with neurons play an essential role in epilepsy ^10-12,42^. The role of astroglial gap junctions in redistributing ions and neurotransmitters in epilepsy models are well investigated ^12,43^. However, the function of astroglial glutamate transporters in epilepsy are less understood. Our results provide direct evidence for the importance of astroglial glutamate transporters for balancing brain excitability. On the one hand, in the absence of glutamate transporter EAAT2a, which is predominantly expressed on astroglia, we observed spontaneous seizures recruiting both neuronal and astroglial networks. In addition, light-induced reflex seizures are apparent in both cellular networks. Such evidence of hyperexcitability is likely due to impaired glutamate clearance in *eaat2a*^-/-^ mutants, leading to a massive transient surge of extracellular glutamate ^31^, as measured by iGluSnFR imaging. The observed upregulation of glutamate transporter *eaat1b* transcripts further supports this hypothesis. On the other hand, we observed that both neuronal and astroglial networks are hypoactive during interictal periods. We argue that this hypoactive state might reflect the reduced availability of glutamate due to depletion of the presynaptic glutamate pool after seizures in combination with impaired glutamate recycling. Since astroglial uptake of glutamate is impaired in *eaat2a*^-/-^ mutants, the conversion of glutamate to glutamine is likely reduced, resulting in an impaired glutamate-glutamine cycle ^44^. This eventually leads to lesser glutamate reconversion in the presynaptic neurons, leading to hypoactivity ^45^. Alternatively, impaired glutamate clearance from the synaptic cleft can lead to desensitization or reduction of postsynaptic glutamate receptors, especially ionotropic AMPA receptors ^46,47^. Consequently, the sensitivity to basal extracellular glutamate fluctuations may be decreased in these animals, and only higher levels of extracellular glutamate induced by sensory stimulation lead to the observed hyperexcitability. A third possibility is that the spontaneous release of glutamate from the presynapse may be suppressed by a negative feedback by neuropeptide modulators. As such, neuropeptide Y, which indirectly inhibits presynaptic glutamate release, has been shown to be upregulated in patients with resistant epilepsy and rodent models ^48,49^. It is likely that all these mechanisms contribute to the switch between the hypoactive inter-ictal state and epileptic seizures. In fact, not only the recurrent spontaneous seizures of *eaat2a*^-/-^ mutants, but also the reduced interictal brain activity mirror observations in human patients; de novo mutations in the human orthologue EAAT2 cause lower residual brain activity and profound intellectual disability ^18,19^.

In recent years, pharmacological and genetic zebrafish models have been used to advance the understanding of epileptogenesis ^28-30,50–57^. Many of these studies focus on the inhibitory system by either pharmacologically or genetically manipulating GABA receptors. However, a better understanding of the major excitatory neurotransmitter system is of great importance, also considering that glutamate is the primary precursor of GABA ^44^. In our model of targeting the glutamate transporter EAAT2a, we found several similarities to existing zebrafish models. Considering seizure propagation, we found that seizures in *eaat2a*^-/-^ mutants initiate in midbrain and hindbrain regions that include important primary processing areas such as the optic tectum (homologous to mammalian superior colliculus ^58^) and the cerebellum ^59^. Interestingly, the anterior forebrain (homologous to mammalian neocortex ^60^) is recruited only with a significant delay. Compared to the PTZ-induced model leading to a lack of neuronal inhibition, our results are similar to findings of one study ^10^, yet contradicting another study showing epileptic propagation from anterior to posterior brain regions ^29^. However, both studies found neuronal microcircuits to be a crucial step in seizure propagation. In line with this idea, we observed not only global but also localized seizures in *eaat2a*^-/-^ mutants, all starting in subcortical regions. These findings support the hypothesis that highly connected hub-like regions play an important role as gate keepers between seizure foci and global brain networks ^61^. Furthermore, it suggests that the recruitment of seizure-prone (ictogenic) hubs during seizure propagation might be more crucial than the initial dysfunction itself. It is likely that this recruitment of specific circuits is dependent on the current brain state, which has been suggested to influence seizure probability ^62^. In fact, *eaat2a*^-/-^ animals suffering global reflex seizures also always exhibited spontaneous global seizures. Hence, future studies in our model may help to understand the transition from local to global seizure networks, and the role of astroglia in these transitions.

We also observed fundamental differences between our *eaat2a*^-/-^ mutants and existing zebrafish models. Our model reflects that epilepsy is more than just a seizure disorder. In fact, subsequent neurobiological and cognitive consequences is part of the epilepsy definition ^9^. While existing zebrafish models helped to observe several aspects of epileptic hyperexcitability ^10,29,30,50^, our *eaat2a*^-/-^ mutants display reduced inter-ictal brain activity. The low neuronal and astroglial network activity in *eaat2a*^-/-^ mutants likely reflects pathological changes in the brain ^40^, potentially corresponding to intellectual disabilities in patients with EAAT2 de novo mutations ^18,19^. We also observed decreased brain size in *eaat2a*^-/-^ larvae, possibly reflecting cerebral atrophy found in patients with EAAT2 mutations ^18,19^. Furthermore, our findings on overall reduced locomotion in *eaat2a*^-/-^ zebrafish is consistent with reduced muscle tone in human patients ^18,19^. Hence, relying solely on increased locomotion as seizure readout in zebrafish models may miss important aspects of epilepsy. Accordingly, focusing not only on the seizure-related hyperactivity but also on the pathophysiology of reduced inter-ictal activity might help unraveling important underlying details of the combined clinical presentation of DEE. Our findings in *eaat2a*^-/-^ mutant zebrafish may also have a broader relevance for epilepsy research. The current clinical dichotomy between focal and generalized seizures is operational and may not reflect mechanistic distinctions ^63–65^. Recent research in human patients using advanced neurophysiological methods and functional imaging is transforming our understanding on ictogenesis. Even the archetypical ‘generalized’ absence seizures involve rather selective parts of the brain, and not the entire cortex as suggested by conventional scalp electroencephalogram ^65,66^. Our transparent zebrafish model enables detailed whole-brain imaging of widespread glia-neuron networks ^67,68^. We observed at high temporal resolution a local subcortical seizure origin and subsequent global propagation. It has been proposed that all seizures in fact initiate within local networks, and subsequent spreading results from a lost balance between local and global network connectivity ^69,70^. Subcortical networks are likely of particular interest, given that they have been shown to be intimately involved in seizures traditionally thought to arise from cortical lobes ^71,72^. Furthermore, the *eaat2a*^-/-^ model enables a certain temporal control of seizure onset through light-induction, providing a prominent window to study ictogenesis. Future studies in our *eaat2a*^-/-^ model may help to improve the understanding on interactions between local and global seizure networks. For all these reasons, we propose to use our novel epilepsy model comprehensively to further the understanding of underlying epileptogenic mechanisms. We argue that our astroglial *eaat2a*^-/-^ mutant zebrafish model will provide an unexplored platform for identifying new treatment approaches, especially taking into account gliotransmission as a promising novel target ^73–76^.

## METHODS

### Fish husbandry

Zebrafish (*Danio rerio*) were kept under standard conditions ^77^. In this study, WIK and Tübingen wildtype strains were used. For calcium imaging experiments, *eaat2a*^+/-^ mutant animals were outcrossed with *Tg(elavl3:GCaMP5G);nacre*^-/-^ ^78^(epifluorescence microscope), *Tg(gfap:Gal4)nw7;Tg(UAS:GCaMP6s)* ^10,79^ fish (two-photon spontaneous recordings) and *Tg(elavl3:GCaMP6s)* ^17^ (two-photon light-stimulation recordings). For experiments, adult *eaat2a*^+/-^ animals were set up pairwise and embryos were raised in E3 medium (5 mM NaCl, 0.17 mM KCl, 0.33 mM CaCl2, 0.33 mM MgSO4, 10^-5^ % methylene blue) (Zürich) or in egg water (60 mg/l marine salt, 10^-4^ % methylene blue) (Trondheim) at 28°C. As control, *eaat2a*^-/-^ larvae were compared to *eaat2a*^+/+^ and *eaat2a*^+/-^ siblings. All experiments were conducted in accordance with local authorities (Zürich Switzerland: Kantonales Veterinäramt TV4206, Trondheim Norway: directive 2010/63/EU of the European Parliament and the Council of the European Union and the Norwegian Food Safety Authorities).

### In situ hybridization

*eaat2a* (ENSDARG00000102453) cloning into the TOPO pCRII vector (TA Cloning Kit Dual Promoter, Invitrogen, Basel, Switzerland) and preparation of digoxigenin (DIG)-labeled antisense RNA probes is described elsewhere ^15,16^. RNA probes were applied on whole-mount zebrafish larvae (5 dpf) and adult brain cross sections at a concentration of 2 ng/μL at 64°C overnight ^80^. Stained and paraformaldehyde (PFA) post-fixed embryos (in glycerol) and sections were imaged with an Olympus BX61 brightfield microscope. Images were adjusted for brightness and contrast using Affinity Photo Version 1.8 and assembled Affinity Designer Version 1.7.

### Immunohistochemistry

Generation of the chicken anti-EAAT2a (zebrafish) antibody is described elsewhere ^16^. 5 dpf larvae were fixed in 4% PFA in PBS (pH 7.4) at room temperature for 40 minutes. Embryos were cryo-protected in 30% sucrose in PBS at 4°C overnight, embedded in Tissue Freezing Medium TFMTM (Electron Microscopy Sciences), cryo-sectioned at 14-16 μm and mounted onto Superfrost slides (Thermo Fisher Scientific). Immunohistochemistry was performed as described before ^16^. Chicken anti-EAAT2a 1:500, mouse anti-synaptic vesicle 2 (IgG1, 1:100, DSHB USA), mouse anti-acetylated tubulin (IgG2b, 1:500, Sigma 7451) and mouse anti-glutamine synthetase (IgG2a, 1:200, EMD Millipore, MAB302) were used as primary antibodies. Secondary antibodies were goat antichicken Alexa Fluor 488, goat anti-mouse IgG2a Alexa Fluor 568, goat anti-mouse IgG2b Alexa Fluor 647 and goat anti-mouse IgG1 Alexa Fluor 647, all 1:500 (all from Invitrogen, Thermo Fisher Scientific). Slides were cover-slipped using Mowiol (Polysciences) containing DABCO (Sigma-Aldrich) and imaged with a TCS LSI confocal microscope (Leica Microsystems). Images were adjusted for brightness and contrast using Affinity Photo Version 1.8 and assembled Affinity Designer Version 1.7.

### CRISPR/Cas9-mediated mutagenesis and genotyping

CRISPR target sites for *eaat2a* (ENSDARG00000102453) of the 5’-GG(N18)NGG-3’ motif favorable for T7-mediated in vitro transcription ^81^ were selected using the https://chopchop.cbu.uib.no/ and www.zifit.partners.org prediction tools. Synthesis of single guide RNA (sgRNA) was performed using a PCR based approach as follows. dsDNA of the target region was amplified with a high fidelity Phusion polymerase (New England Bio Labs) using the forward primer sg1 together with the common reverse primer sg2 (Supplementary Table 1). sgRNA was T7 in vitro transcribed (MEGAshortscriptTM T7 Transcription Kit, Ambion) and subsequently purified using the MEGAclearTM Kit (Ambion).

The injection mix consisting of 160 ng/μL sgRNA, 1 μg/μL Cas 9 protein (Flag/ or GFP-tagged Cas9 kindly provided by Prof. Dr. C. Mosimann and Prof. Dr. M. Jineck) and 300 mM KCl was incubated at 37°C for 10 minutes to enable Cas9/sgRNA complex formation. One-cell staged embryos were injected with 1 nL injection mix into the cell. Mutation rate efficiency was tested by genotyping a pool of around 10-15 injected F0 larvae per clutch while their siblings were raised to adulthood. Adult F0 crispant fish were outcrossed to Tübingen wild-type animals. F1 embryos were genotyped at 3 dpf by larval tail biopsies ^82^ and raised in single tanks. The *eaat2a* target site was PCR amplified with a fast-cycling polymerase (KAPA2G Fast HotStart PCR kit, KAPA Biosystems) (primers: sense (fw) and antisense (rev, Supplementary Table 1)). Amplicons were cloned into pCR 2.1-TOPO vectors (Invitrogen) and sequenced. The resulting heterozygous mutant line carrying one copy of a −13 null allele was repeatedly outcrossed to wild-type and transgenic fish to generate stable heterozygous F2, F3 and F4 generations. *eaat2a* mutant fish can be genotyped by the above-mentioned PCR amplification and a subsequent gel-electrophoresis, which allows detection of the 13 base pair deletion. Larvae were genotyped after each experiment. After two-photon microscopy, all larvae were genotyped by means of a PCR melting curve analysis using SYBR Green (PowerUp SYBR Green Master Mix, Thermo Fisher Scientific). For qRT-PCR experiments, wild-type progeny of *eaat2a*^+/-^ incrosses were selected by pre-genotyping using the Zebrafish Embryo Genotyper (ZEG, wFluidx) as described previously ^83^. Briefly, 3-4 dpf embryos were loaded individually onto the ZEG chip in 12 μL E3 and vibrated for 10 minutes at 1.4 Volts. Subsequently, 8 μL of each sample E3 was used directly for PCR as described above (KAPA Biosystems). Embryos were kept in 48-well plates until genotyping was achieved.

### Zebrafish survival analysis and length measurements

Embryos were raised separately in 24-well plates containing 1.5 mL E3 medium, which was changed daily. From 5 dpf on, larvae were fed manually. Animal survival was monitored daily over an 11-day period. From 1 to 7 dpf, larvae were anaesthetized by tricaine (0.2 mg/mL) and individually imaged with an Olympus MVX10 microscope. Body lengths were measured along the spinal cord using Fiji ImageJ (National Institutes of Health) and further analyzed using R software version 3.6.0 with the RStudio version 1.2.1335 interface ^84,85^.

### Behavioral analysis

Swimming patterns of 5 dpf larvae were recorded using the ZebraBox system (ViewPoint Life Sciences). The room was kept at 27°C throughout the recordings. Larvae were individually placed at randomized positions of 48-well plates containing 1 mL E3 medium and transferred into the recording chamber for a minimum of 10 minutes acclimatization. Subsequently, recordings of 30 minutes in normal light conditions (20% light intensity) were performed. An automated camera tracked individual larvae (threshold black 30) and detected the distance and duration moved by each larva exceeding 1 mm per second in accumulating five-second intervals. Data processing and analysis was done with a custom code written in R with the RStudio interface ^84,85^. Software-related artefacts were removed in a blinded manner. Velocity of long-lasting bursts were calculated as distance divided by time for every five-second interval in which the animal moved at least 3.5 seconds.

### Combined LFP experiments together with epifluorescence iGluSnFR imaging

Simultaneous LFP (local field potential) recordings of seizure activity and epifluorescence imaging of *iGluSnFR* signals were performed in 5 dpf *Tg(gfap:iGluSnFR)* zebrafish larvae ^10^. First, zebrafish larvae were anesthetized in 0.02% MS222 and paralyzed by α-bungarotoxin injection ^86^. Next, the larvae were embedded in 1-1.5% low melting point agarose (Fisher Scientific) in a recording chamber (Fluorodish, World Precision Instruments). Artificial fish water (AFW, 1.2 g marine salt in 20 L RO water) was constantly perfused during the experiments. For LFP recordings, a borosilicate glass patch clamp pipette (9-15 MOhms) loaded with teleost artificial cerebrospinal fluid ^87^ (ACSF, containing in mM: 123.9 NaCl, 22 D-glucose, 2 KCl, 1.6 MgSO_4_ 7H_2_O, 1.3 KH_2_PO_4_, 24 NaHCO_3_, 2 CaCl_2_ 2H_2_O) was inserted in the forebrain ^75^. LFP recordings were performed by a MultiClamp 700B amplifier, in current clamp mode at 10 kHz sampling rate, and band pass filtered at 0.1–1000 Hz. For imaging *iGluSnFR* signals, microscopy images were collected at 10Hz sampling rate, using a Teledyne QImaging QI825 camera, in combination with Olympus BX51 fluorescence microscope and Olympus UMPLANFL 10X water immersion objective. Data acquisition of LFP signals were done in MATLAB (Mathworks), and *iGluSnFR* signals were done in Ocular Image Acquisition Software. All data were analyzed using MATLAB.

### Quantitative reverse transcription PCR

5 dpf larvae were anesthetized on ice and brains dissected using an insect pin and a syringe needle in a dish containing RNA later (Sigma-Aldrich). Total RNA of equal pools of larval brains was extracted using the ReliaPrep kit (Promega). RNA was reverse transcribed to cDNA using the Super Script III First-strand synthesis system (Invitrogen) using 1: 1 ratio of random hexamers and oligo (dt) primers.

The qPCR reactions were performed using SsoAdvanced Universal SYBR Green Supermix on a CFX96 Touch Real-Time PCR Detection System (Bio-rad). Primer (Supplementary Table 2) efficiencies were calculated by carrying out a dilution series. After the primer efficiencies were determined equal, brain samples were used for qPCR using 10 ng of cDNA per reaction. The controls “no reverse transcription control” (nRT control) and “no template control” (NTC) were performed with every qPCR reaction. *g6pd* and *b2m* were chosen as reference genes. All reactions were performed in technical triplicates. Data was analyzed in CFX Maestro Software from Bio-Rad and Microsoft Excel. Statistical analysis was performed between dCt values.

### Calcium imaging and data analysis

#### a. Epifluorescence microscope

5 dpf larvae in the *Tg(elavl3:GCaMP5G)* background were individually immobilized in a drop of 1.8% NuSieveTM GTGTM low melting temperature agarose (Lonza) with the tail freed from agarose in a small cell culture dish (Corning Incorporated) filled with water. Calcium signals were recorded using an Olympus BX51 WI epifluorescence microscope and by means of a camera (4 Hz sampling rate) and the VisiView software established by the Visitron Systems GmbH.

In each experiment, the GCaMP5G fluorescence signal of manually selected ROIs (whole brain, forebrain, midbrain and hindbrain) was extracted using Fiji ImageJ (National Institutes of Health). For each time point, the mean intensity of each ROI was measured and further processed using a custom script in R with the RStudio interface^84,85^. The baseline (F_0_) of every ROI was calculated as 1^st^ percentile of the entire fluorescence trace per fish. Subsequently, the fractional change in fluorescence (ΔF/F_0_) of each ROI was computed to normalize the values obtained. In order to compare basal activity between *eaat2a*^-/-^, *eaat2a*^+/-^ and *eaat2a*^+/+^ larvae, the standard deviation of F0 for the whole brain was calculated and averaged over five two-minute time windows per animal (same random windows for all fish, but adjusted if during seizure). Seizure duration in *eaat2a*^-/-^ mutants was defined as the period from first time point where ΔF/F is greater than 50% (seizure initiation) until first time point where ΔF/F is below 50% in the midbrain.

#### b. Two-photon microscope

Two-photon calcium recordings were performed on 5 dpf *eaat2a* mutant *Tg(gfap:Gal4)nw7;Tg(UAS:GCaMP6s)* zebrafish^10^. Larvae were pre-selected based on morphological phenotype, paralyzed by injecting 1 nL of α-bungarotoxin (Invitrogen BI601, 1 mg/mL) into the spinal cord, embedded in 1.5-2% low melting point agarose (LMP, Fisher Scientific) in a recording chamber (Fluorodish, World Precision Instruments) and constantly perfused with artificial fish water (AFW, 60 mg/l marine salt in RO water) during imaging. The cerebral blood flow was assessed in the prosencephalic arteries or anterior cerebral veins^88^ before and after the experiment, and animals without cerebral blood flow were excluded from the final analysis. Imaging was performed in a two-photon microscope (Thorlabs Inc and Scientifica Inc) with a x16 water immersion objective (Nikon, NA 0.8, LWD 3.0, plan). Excitation was achieved by a mode-locked Ti:Sapphire laser (MaiTai Spectra-Physics) tuned to 920 nm. Single plane recordings of 1536 x 650 pixels were obtained at an acquisition rate of 24 Hz. The plane was oriented towards the glial cells in the boundary region between the telencephalon (anterior forebrain) and the anterior thalamus (posterior forebrain). First, spontaneous calcium activity was measured for 60 minutes. Subsequently for a subgroup of the fish, a custom-made Arduino was used to apply two 10-second red-light stimuli (625 nm). Light was flashed after 5 and 10 minutes of the total duration of 15 minutes.

Images were aligned, cells detected and ΔF/F0 relative to the baseline calculated using the algorithm previously described^89,90^, with adaptions as follows. Glial cells along the ventricle were semi-automatically detected and subsequently assigned manually according to their location^10,89,91^. Baseline was computed within a moving window of 80 seconds as the 8^th^ percentile of activity defining noise with a Gaussian curve fit using an adapted algorithm^92,93^. Resampled ΔF/F0 traces of 4 Hz were used to detect glial calcium signals. Calcium events were identified by detecting events significantly different from noise level within a 95% confidence interval^92,93^. For spontaneous recordings, glial activity was quantified in the following two-minute time windows: basal (between seizure activity, averaged over six time points per fish), preictal (preceding seizure onset) and ictal (during seizure). Seizure onset frames were defined as time points where the averaged ΔF/F0 of all cells not located along the ventricles reached 50. For light stimuli recordings, glial activity was compared between one minute prior to and one minute following light stimuli. In each period, a cell was considered active if at least one event occurred. The overall activity of active cells was calculated by the trapezoidal numerical integration method (function *trapz*, MATLAB) to get the area under the curve (AUC). The amplitude of each event was determined as its respective maximum peak.

Light stimulus assay of neuronal activity was performed on 5 dpf *eaat2a*^+/-^ *Tg(elavl3:GCaMP6s)* incross progenies. Mutant larvae were pre-selected, paralyzed and embedded as described above. After the agarose solidified for 10 minutes, AFW was added as an immersion medium before 20 more minutes of agarose settling. The animals were acclimatized in the setup for 20 minutes before volumetric recordings of ten planes of 1536 x 650 pixels were obtained using a Piezo element at a rate of 2,43 Hz. Excitation was achieved as described above. Baseline spontaneous calcium activity was recorded for ten minutes in darkness, followed by a red-light stimulus train using a red LED (LZ1-00R105, LedEngin; 625-nm) placed in front of the animal. Five light stimuli with five-minute inter-stimulus intervals were applied by an Arduino-device. Images were aligned as described above. Regions of interest (ROIs) were manually drawn on one single plane of maximal information. F0 baseline was computed as 1^st^ percentile of the entire fluorescence trace per ROI.

### Statistical analysis

Statistical analysis was done using R software version 3.6.0 with the RStudio version 1.2.1335 interface^84,85^. Data sets were tested for normality using quantile-quantile plots and Shapiro-Wilk test. Normally distributed data was analyzed using Welch two-sample unpaired t-test (Fig. 5i and m). Wilcoxon rank-sum test was used for non-paired analysis (Fig. 2g, Fig. 5g, h, j-l), Wilcoxon signed rank test for paired analysis (Fig. 5e, f) and two-sample Kolmogorov-Smirnov test for equality between distributions (Fig. 3f). Kruskal-Wallis rank-sum test with Wilcoxon rank-sum posthoc test was used for nonpaired analysis between all three genotypes (Fig. 2b, Fig. 3c, Fig. 4j). *P* <0.05 was considered as statistically significant.

### Data and Code availability

The datasets and codes supporting this study have not been deposited in a public repository, but are available from the corresponding authors upon request.

## ACKNOWLEDGEMENTS

We thank R. MacDonald and W. Harris (Cambridge University, UK), A. Schier and F. Engert (Harvard University, USA), M. Ahrens (HHMI, Janelia Farm, USA) and K. Kawakami (SOKENDAI, The Graduate University for Advanced Studies, Japan) for transgenic lines. We thank C. Mosimann and M. Jinek for kindly providing us with Cas9 protein. We thank M. Walther, K. Dannenhauer and H. Möckel in Zürich, and S. Eggen, V. Nguyen, M. Andresen and the zebrafish facility support team in Trondheim for excellent technical and animal support. We also thank E. Brodtkorb (St. Olav’s University Hospital and NTNU, Norway) and M. Gesemann for critical comments on the manuscript and the Yaksi and Neuhauss lab members for stimulating discussions. This work was funded by the Swiss National Science Foundation Grant 31003A_173083 (A.L.H., N.R., S.N., S.C.F.N), UZH Forschungskredit Candoc Grant K-74417-01-01 (A.L.H.), Flanders Science Foundation (FWO) Grant (E.Y.), RCN FRIPRO Research Grant 314212 (E.Y.), RCN FRIPRO Research Grant 314189 (N.J.Y.), Medical Student’s Research Programme NTNU (A.J.), and The Liaison Committee for Education, Research and Innovation in Central Norway (‘Samarbeidsorganet’) Grant (S.M-S., N.J-Y., E.Y.). Work in the E.Y. lab is funded by the Kavli Institute for Systems Neuroscience at NTNU.

## AUTHOR CONTRIBUTIONS

A.L.H., N.J.Y., E.Y. and S.C.F.N conceptualized the study; A.L.H., A.J., N.R., S.N. and E.A. performed experiments; A.L.H., A.J., N.R., E.A., N.J.Y. and E.Y. analysed data; A.L.H., A.J., L.L., N.J.Y. and E.Y. developed custom R and Matlab codes; A.L.H., N.J.Y. and E.Y. prepared all figures; A.L.H., N.J.Y., E.Y. and S.C.F.N wrote the manuscript; A.L.H., A.J., S.M.S., N.J.Y., E.Y. and S.C.F.N. edited the manuscript with the help of all authors; N.J.Y., E.Y and S.C.F.N. acquired funding and supervised the students.

## ADDITIONAL INFORMATION

Correspondence and requests for materials should be addressed to N.J.Y., E.Y. or S.C.F.N.

