## Supplementary information for "Loss of glutamate transporter *eaat2a* leads to aberrant neuronal excitability, recurrent epileptic seizures and hypoactivity"

### Supplementary figures

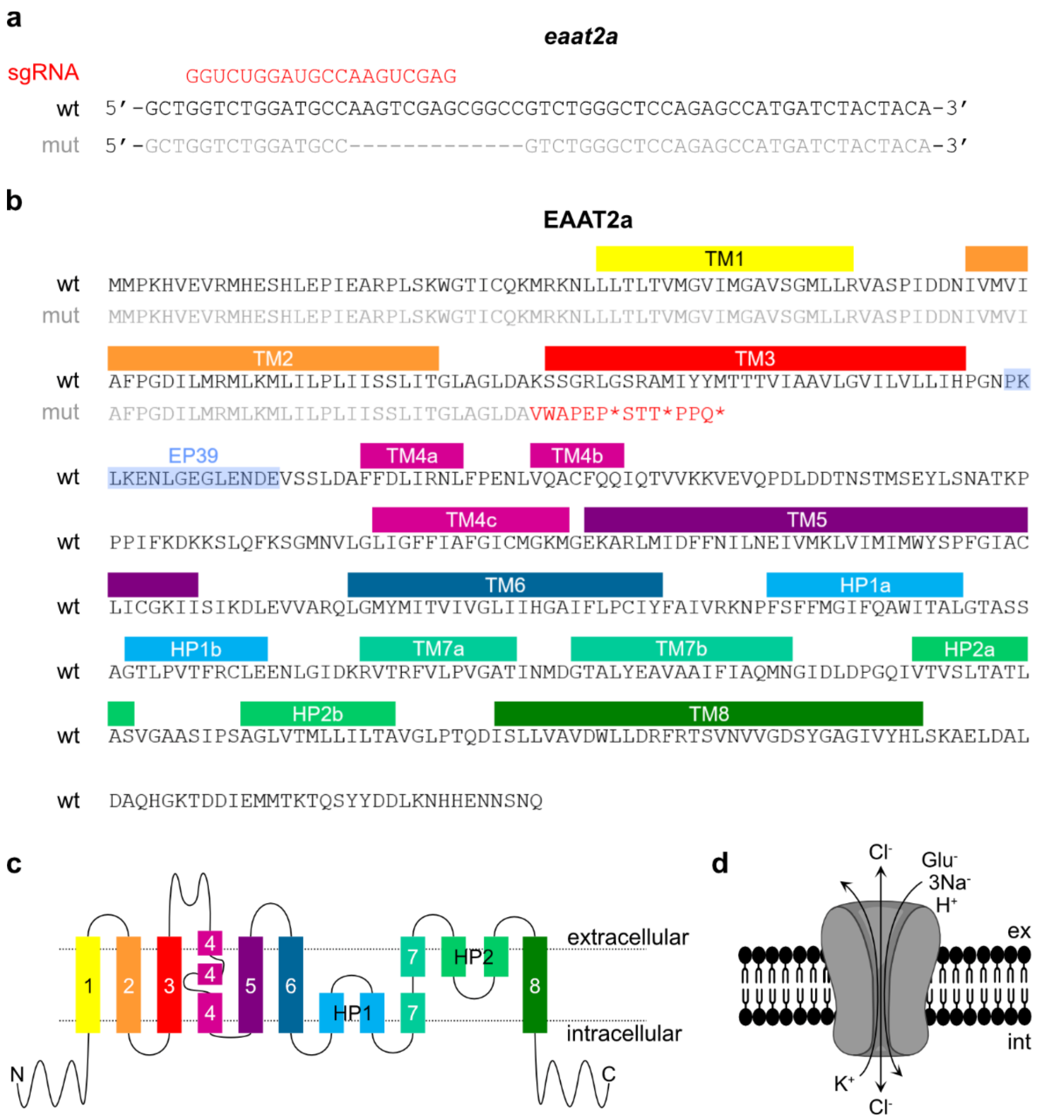


**Supplementary Figure 1: Wild-type and mutant *eaat2a* sequence. a** Genomic DNA sequence of wild-type (black font) and mutant (gray font) *eaat2a* harboring a -13 base pair deletion. Target sequence of the single guide RNA used is represented in red font. **b** EAAT2a wild-type (black font) and mutant (gray font) protein sequence with corresponding transmembrane domains (TMs) and hairpins (HPs) labelled. The -13 bp mutation leads to three premature stop codons (*) within TM3 at amino acids 109, 113 and 117. EP39 (epitope 39) highlights the anti-EAAT2a antibody recognition site. **c** Two-dimensional schematic representation of EAAT2a protein structure. Color code for TM and HP labelling corresponds to color code in **b**. **d** Stoichiometry of EAATs.

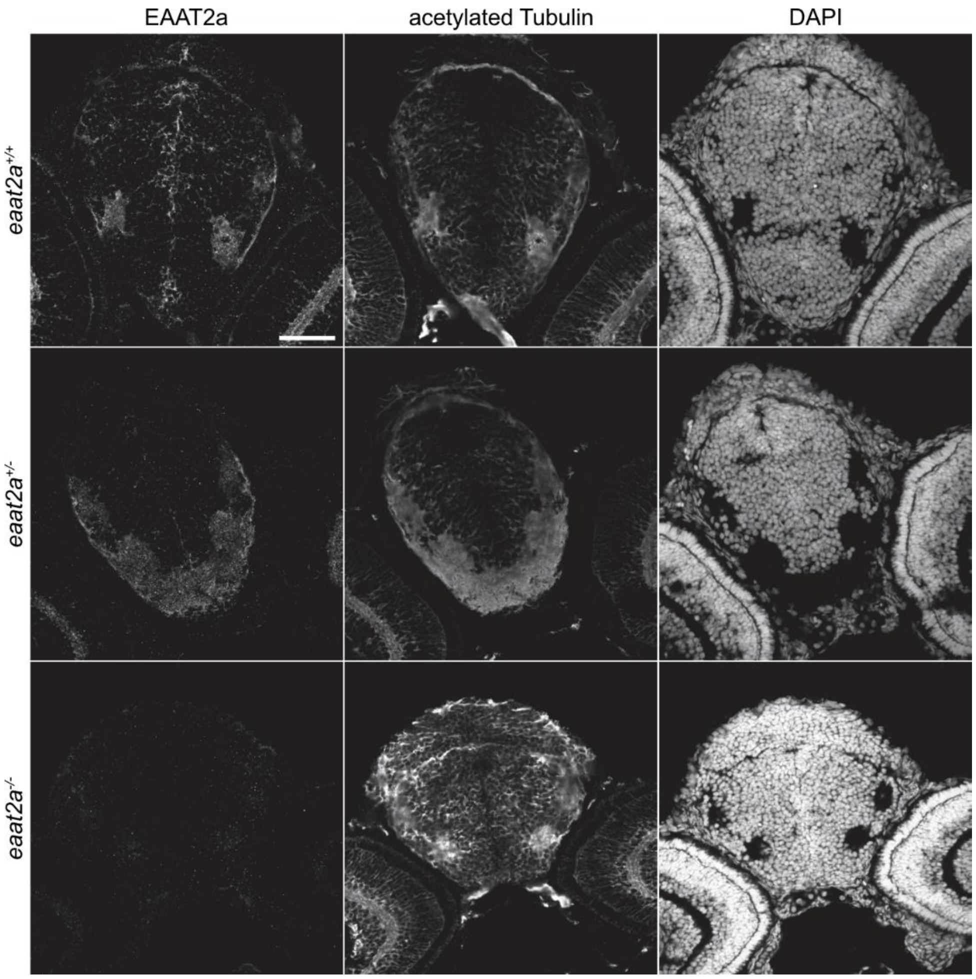


**Supplementary Figure 2: Confirmation of EAAT2a protein knockout.** Immunostaining of EAAT2a and acetylated tubulin (acT) together with DAPI on 3 dpf *eaat2a*$^{+/+}$ (top row), *eaat2a*$^{+/-}$ (middle row) and *eaat2a*$^{-/-}$ (bottom row) larvae. Scale bar is 50 µm.

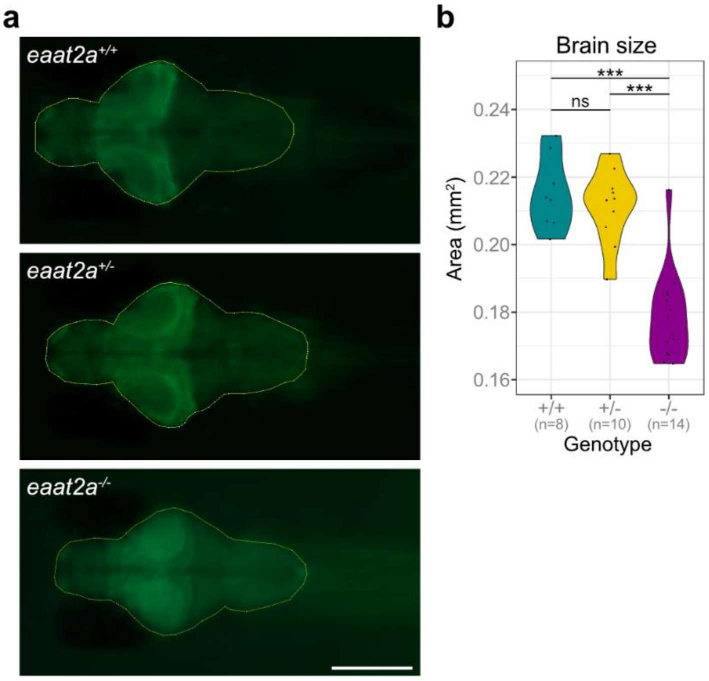


**Supplementary Figure 3: Overall brain size in 5dpf *eaat2a*$^{-/-}$ mutants is decreased. a** Representative wide-field microscopy images of *eaat2a*$^{+/+}$ (top), *eaat2a*$^{+/-}$ (middle) and *eaat2a*$^{-/-}$ (bottom) larvae. Yellow selection shows region of interest used to measure total brain area by Fiji ImageJ (National Institutes of Health). Scale bar is 250 µm. **b** Brain area compared among genotype. Significance levels: ***p = < 0.001, ns = not significant ($p$ > 0.05), Kruskal Wallis test with Wilcoxon rank-sum posthoc test.

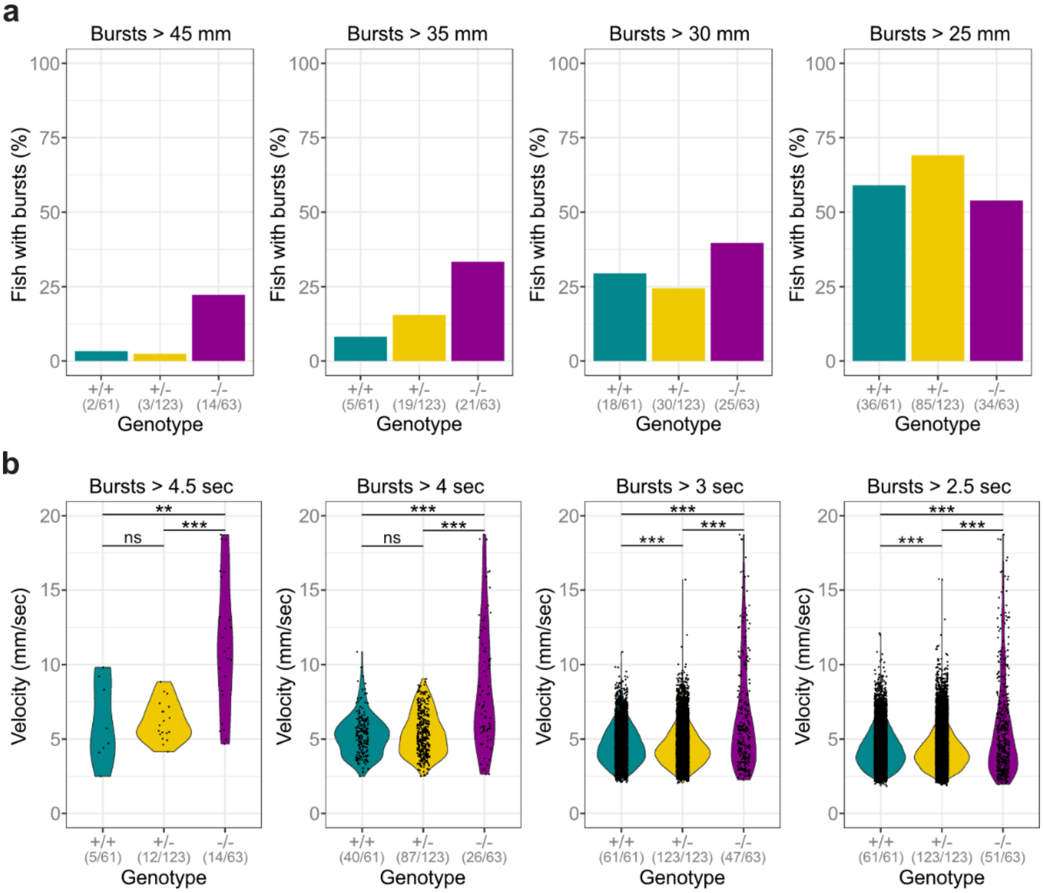


**Supplementary Figure 4: Swim bursts per five-second integral in 5 dpf *eaat2a*$^{-/-}$ larvae are more frequent at higher thresholds**. **a** Proportion of fish showing one or more bursts bigger than threshold indicated in headings (mm). **b** Velocity of all bursts lasting longer than threshold indicated in headings (sec). Significance levels: ***$p$ = < 0.001, **$p$ = < 0.01, ns = not significant ($p$ > 0.05), two-sample Kolmogorov-Smirnov test.

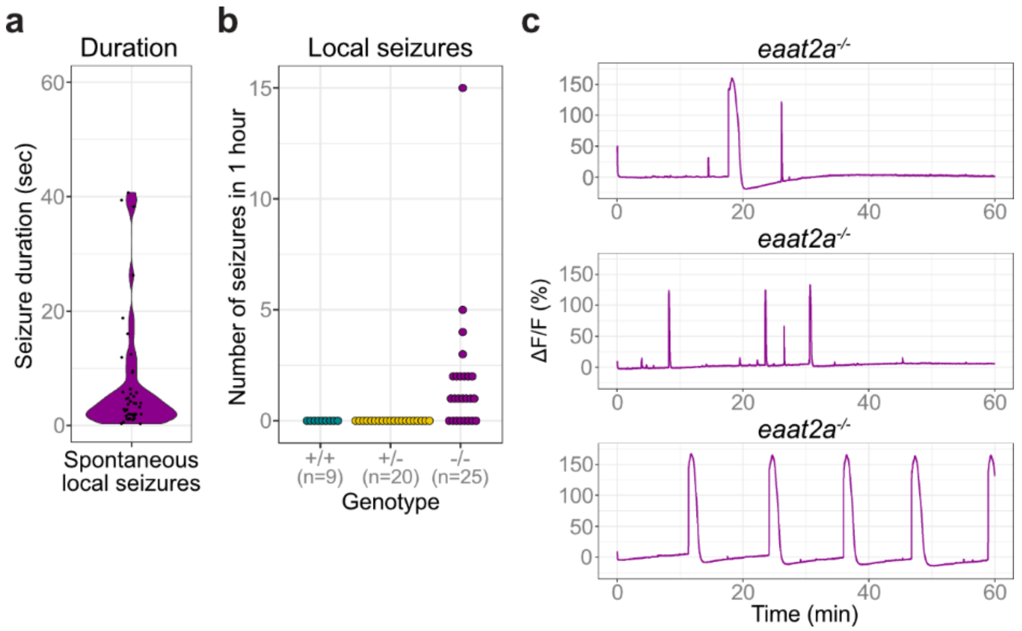


**Supplementary Figure 5: 5 dpf *eaat2a*$^{-/-}$ mutants show not only global but also local seizures not recruiting neurons of the anterior forebrain and lasting less than a minute. a** Duration of local seizures present in *eaat2a*$^{-/-}$ mutants (median 3.4 sec (std ± 10.3 sec)). **b** Number of spontaneous local seizures per animal during 60 minutes. **c** Change in GCaMP5G fluorescence (ΔF/F) over time recorded across the entire brain of three representative *eaat2a*$^{-/-}$ mutant larvae displaying local and global (top), exclusively local (middle) or exclusively global (bottom) spontaneous seizures.

### Supplementary tables

| Primer name | Primer sequence (5' – 3') | Amplicon (bp) |
|---|---|---|
| *eaat2a* sgRNA sg1 | GAAATTAATACGACTCACTATA**GGTCTGGATGCCAAGTCGAG** GTTTTAGAGCTAGAAATAGC | 20 |
| *eaat2a* sgRNA sg2 | AAAAGCACCGACTCGGTGCCACTTTTTCAAGTTGATAACGGACT AGCCTTATTTTAACTTGCTATTTCTAGCTCTAAAAC | |
| *eaat2a* sense fw genotyping | GATGCAGTCGTATGGGAA | 204 |
| *eaat2a* antisense rev genotyping | CCTTCTCCCAGATTCTCC | |

**Supplementary Table 1: List of primers used for CRISPR sgRNA synthesis and target region amplification.**

| Gene | www.ensembl.org/index.html | Fw primer (5' – 3') | Rev primer (5' – 3') | Amplicon (bp) |
|---|---|---|---|---|
| *g6pd* | ENSDARG00000071065 | CTGGACCTGACCTACCATAGCAG | AGGCTTCCCTCAACTCATCACTG | 127 |
| *b2m* | ENSDARG00000053136 | GACAAAGAAGTTTAGTTGGGAGCC | GAATCCATCGCTCCATCG | 76 |
| *eaat1b* | ENSDARG00000043148 | CTGGTTCAAGCCTGCACTCAAC | CTCCTGCGTAGCGTTCGTC | 111 |
| *eaat2b* | ENSDARG00000052138 | GGAATCGAACTTGATCCTGGTC | ATGTCTTGAGTCGGCAGTCC | 143 |
| *mglur3* | ENSDARG00000031712 | GTCTTGATGGCAGGAAACTCTACC | CCACTCCATCTCCATACGCATC | 117 |
| *mglur5b* | ENSDARG00000102067 | CCTGGTCGGAGAGTTTCTGCTG | GCAGACTGCAGCTTTATGGTGATC | 114 |
| *gad1b* | ENSDARG00000027419 | ACAACAATGAGCGCTTCACG | TGAGGATCTCCACCACTTCGAG | 126 |
| *gabra1* | ENSDARG00000068989 | GGTCACCAATGCTCTGGCTG | GAGACCAGGCCTAAGACGATTG | 130 |

**Supplementary Table 2: List of primers used for qRT-PCR experiments**
